# Integrative analysis of papain-like cysteine proteases and cystatins reveals stress-dependent regulatory modules in *Arabidopsis thaliana*

**DOI:** 10.64898/2025.12.31.697236

**Authors:** Shuchi Wu, Xing Yi, Song Li, Bingyu Zhao

## Abstract

Plant papain-like cysteine proteases (PLCPs) and cystatins constitute a major protease-inhibitor system that contributes to plant signaling and responses to biotic and abiotic stress. Although the Arabidopsis genome encodes 31 predicted PLCPs and 7 cystatins, their coordinated regulation has not been systematically evaluated across major stress conditions. Here, we reprocessed and integrated AtGenExpress microarray datasets to analyze the expression patterns of 28 PLCPs and seven cystatins in response to bacterial infection (virulent and avirulent *Pseudomonas syringae*), wounding, and drought stress. In parallel, we performed in vitro enzyme activity assays to assess inhibition specificity of seven Arabidopsis cystatins against five abundantly expressed PLCPs. Finally, we used the support vector machine (SVM) package e1071 in R to integrate co-expression and inhibition data and to generate a hypothesis-generating PLCP-cystatin interaction network. Together, these analyses suggest that Arabidopsis deploys distinct PLCP and cystatin gene subsets in response to virulent versus avirulent bacterial infection, and that transcriptional modules associated with bacterial infection and wounding are largely distinct. In contrast, several drought-associated PLCP-cystatin modules overlap with those associated with basal defense, suggesting partially shared regulatory programs across abiotic and biotic stress.

## Introduction

Proteases are ubiquitous in all organisms and play key roles in regulating diverse biological processes (Supuran, Scozzafava et al. 2002; Haq, Atif et al. 2004; Valueva and Mosolov 2004; Habib and Fazili 2007; Van der Hoorn 2008). In plant and animal genomes, approximately two percent of genes encode proteases (Rawlings, Tolle et al. 2004). Beyond bulk protein turnover, proteases function as regulators of signaling pathways that control development and stress responses, including embryogenesis, flowering, senescence, and immunity (Reeves, Murtas et al. 2002; Bozhkov, Suarez et al. 2005; Rooney, Van’t Klooster et al. 2005; Diaz-Mendoza, Velasco-Arroyo et al. 2014). Plant proteases are commonly grouped by catalytic mechanism into serine, cysteine, threonine, aspartate, and metalloproteases (Beers, Woffenden et al. 2000; van der Hoorn, Leeuwenburgh et al. 2004; Van der Hoorn 2008). Among these, serine and cysteine proteases have been intensively studied, and many cysteine proteases have documented roles in plant immunity (van der Hoorn, Leeuwenburgh et al. 2004; Van der Hoorn 2008). Cysteine proteases in family C1A, also known as papain-like cysteine proteases (PLCPs), are prominent components of programmed cell death (PCD) and immune responses (Rawlings, Tolle et al. 2004; Van der Hoorn 2008). Because uncontrolled PLCP activity can damage host tissues, plants have evolved multiple mechanisms to restrict protease activation and activity (Doehlemann and Hemetsberger 2013). PLCPs are synthesized as inactive zymogens containing N-terminal autoinhibitory prodomains, which are removed during maturation to produce active enzymes (Yamada, Matsushima et al. 2001; van der Hoorn, Leeuwenburgh et al. 2004). Activation is often promoted by acidic pH and can be influenced by elicitors such as H_2_O_2_ or Sodium dodecyl sulfate (SDS) (Yamada, Ohta et al. 1998; Solomon, Belenghi et al. 1999). In addition to intrinsic autoinhibition, plants encode specialized protease inhibitors that suppress PLCP activity at the protein level. Among these, cystatins are competitive inhibitors of PLCPs and are conserved across eukaryotes (Habib and Fazili 2007; Benchabane, Schluter et al. 2010). Plant cystatins typically contain the conserved QxVxG motif required for protease binding, along with conserved residues in the N- and C-termini (Turk and Bode 1991; Benchabane, Schluter et al. 2010).

Land plants employ at least two interconnected layers of immunity. First, pattern recognition receptors detect pathogen-associated molecular patterns (PAMPs) and activate PAMP-triggered immunity (PTI). Second, resistance (R) proteins recognize specific pathogen effectors and activate effector-triggered immunity (ETI), a stronger response that often includes hypersensitive response-associated PCD at infection sites (Jones and Dangl 2006). Multiple PLCPs participate in immune signaling. For example, tomato RCR3 is required for Cf2-mediated resistance to *Cladosporium fulvum* (Rooney, Van’t Klooster et al. 2005). Arabidopsis RD19A is induced during bacterial wilt caused by *Ralstonia solanacearum* and interacts with the effector PopP2 (Bernoux, Timmers et al. 2008). Arabidopsis RD21A also contributes to resistance against the necrotrophic pathogen *Botrytis cinerea* (Shindo, Misas-Villamil et al. 2012). In contrast, several cystatins can negatively modulate immunity by inhibiting host PLCPs. For instance, maize cystatin CC9 is induced during compatible interactions with *Ustilago maydis*; CC9 overexpression suppresses PLCP activity and salicylic acid-responsive PR gene expression, increasing susceptibility (van der Linde, Hemetsberger et al. 2012). Arabidopsis AtCYS1 suppresses pathogen- and wound-triggered PCD (Belenghi, Acconcia et al. 2003). Together, these observations highlight the importance of PLCP-cystatin interplay in regulating immune-associated proteolysis.

PLCPs and cystatins also participate in responses to drought stress. Among Arabidopsis PLCP genes, several were originally identified as responsive to desiccation (Yamaguchi-Shinozaki, Koizumi et al. 1992). RD21A, one of the most abundant drought-induced cysteine proteases, is strongly induced during dehydration (Koizumi, Yamaguchi-Shinozaki et al. 1993; Seki, Narusaka et al. 2002) and has been linked to ABA-associated drought tolerance pathways (Kim and Kim 2013). RD21A contribute to drought-induced resistance to *Pseudomonas syringae* in Arabidopsis (Liu et al. 2020). Drought-induced cysteine proteases have also been described in tomato and wheat (Harrak, Azelmat et al. 2001; Simova-Stoilova, Vaseva et al. 2010). Cystatins can influence drought tolerance as well; for example, ectopic expression of rice oryzacystatin-I enhances drought tolerance in soybean and Arabidopsis (Quain, Makgopa et al. 2014), while several maize cystatins are downregulated during water starvation (Massonneau, Condamine et al. 2005). Because drought stress and bacterial infection can both modulate stomatal behavior, plants may deploy partially overlapping regulatory pathways to control stomatal closure in response to abiotic and biotic cues (Melotto, Underwood et al. 2006; Atkinson and Urwin 2012). However, the extent to which PLCP-cystatin modules are shared across these stresses has not been comprehensively assessed.

Co-expression network analysis is widely used to associate genes of unknown function with biological pathways and regulatory programs (Atias, Chor et al. 2009). The approach is based on the premise that genes involved in a shared pathway, regulated by common factors, or participating in the same functional complex often exhibit similar expression patterns across conditions. Co-expression analyses have helped identify genes involved in diverse processes, including metabolic pathways, circadian regulation, and pathogen resistance (Atias, Chor et al. 2009; Leal, Perez et al. 2013). Public microarray resources such as AtGenExpress provide a standardized platform for exploring coordinated expression patterns across tissues and stresses in Arabidopsis.

Support vector machines (SVMs) are supervised learning models originally developed for binary classification (Cortes and Vapnik 1995). SVMs have been applied to high-dimensional gene expression datasets to classify samples and infer functional associations (Brown, Grundy et al. 2000). In the context of PLCP-cystatin analysis, experimentally supported protease-inhibitor relationships can be used as training examples to integrate multiple co-expression features and generate a hypothesis-generating network of candidate associations.

Despite extensive work on individual PLCPs and cystatins, studies linking PLCP-associated immunity and abiotic stress responses remain fragmented. A systematic analysis of PLCP and cystatin expression across major stresses, combined with experimental assessment of inhibition specificity, can provide a useful resource for prioritizing candidate protease-inhibitor modules for functional testing.

In this study, we analyzed co-expression patterns of 28 PLCPs and seven cystatins in Arabidopsis in response to biotic and abiotic stress by reprocessing and integrating AtGenExpress microarray datasets. For biotic stress, we analyzed expression profiles from wild-type (Col-0) plants inoculated with virulent *Pseudomonas syringae* ES4326 or the same strain carrying AvrRpt2 (TAIR Accession ID: 1008031517) (Wang, Weaver et al. 2005). For abiotic stress, we analyzed wounding and drought time-course datasets (TAIR Accession IDs: 1007966439 and 1007966668) (Kilian, Whitehead et al. 2007). In parallel, we performed in vitro protease activity and inhibition assays for five abundantly expressed PLCPs and seven cystatins. Finally, we integrated co-expression features and inhibition results using the SVM package e1071 in R (v3.2.0) (Meyer and Wien 2014) to generate a hypothesis-generating PLCP-cystatin association network.

## Results and Discussion

### Expression Profiles of Seven Cystatins and Five PLCPs of Arabidopsis Infected with *Pseudomonas syringae* ES4326

Several Arabidopsis PLCP and cystatin genes were reported to involve in plant immunity (Rawlings, Tolle et al. 2004, Habib and Fazili 2007, Van der Hoorn 2008). In this study, we characterized the expression patterns of seven Arabidopsis cystatins and five PLCPs under the condition with pathogen inoculation. The five PLCPs were selected because of their relatively higher expression levels either in leaf or root tissue (Supplementary Information Figure. S1). The gene expression patterns were generated from the AtGenExpress microarray data of Arabidopsis thaliana (Col-0) plants challenged with *Pseudomonas syringae* strain (*P. syringae* ES4326), with or without the avirulence effector AvrRpt2 (TAIR Accession ID: 1008031517). Figure 1a &b summarizes the gene expression patterns of the selected genes in the condition with virulent or avirulent pathogen inoculation.

**Figure 1.**
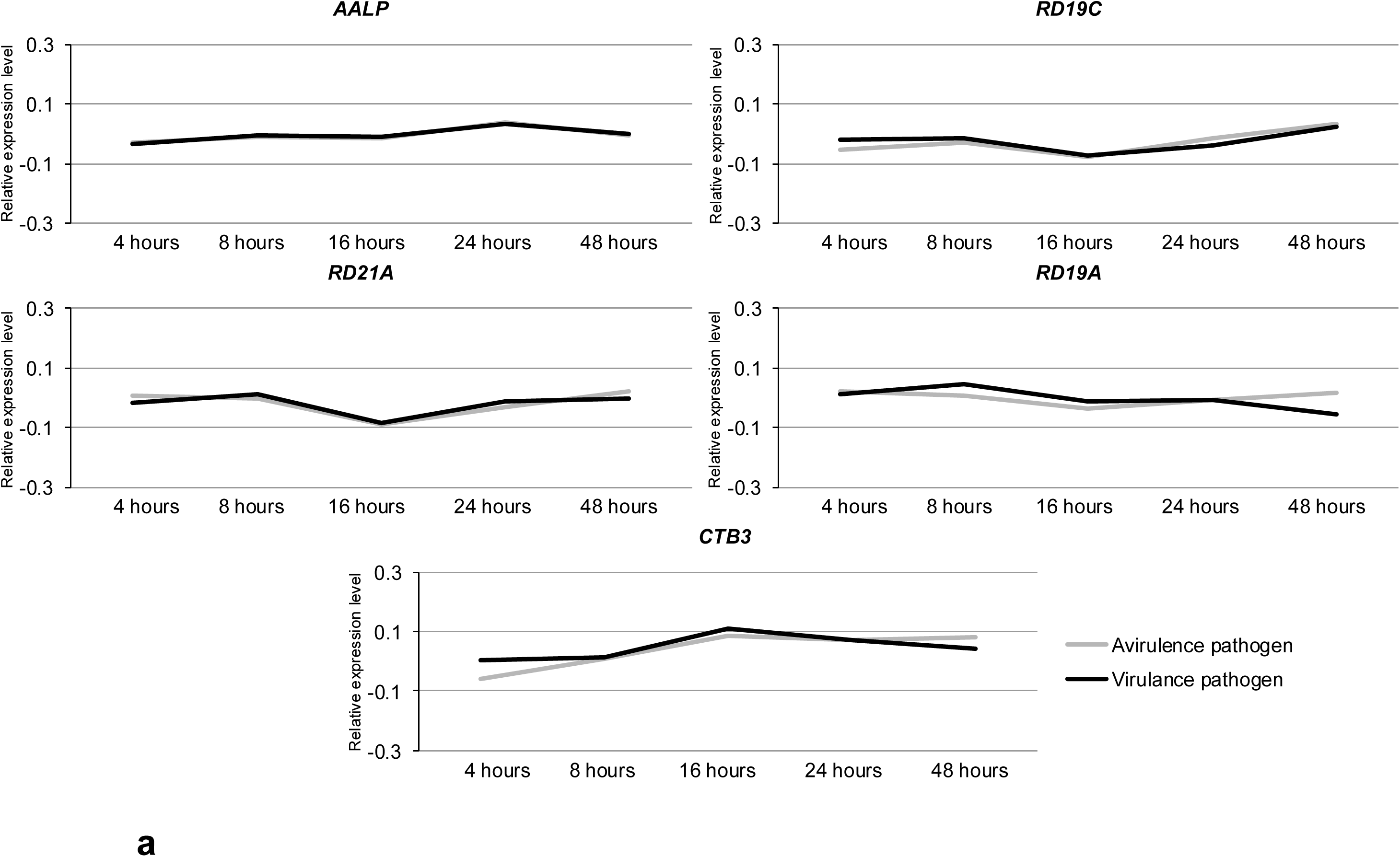
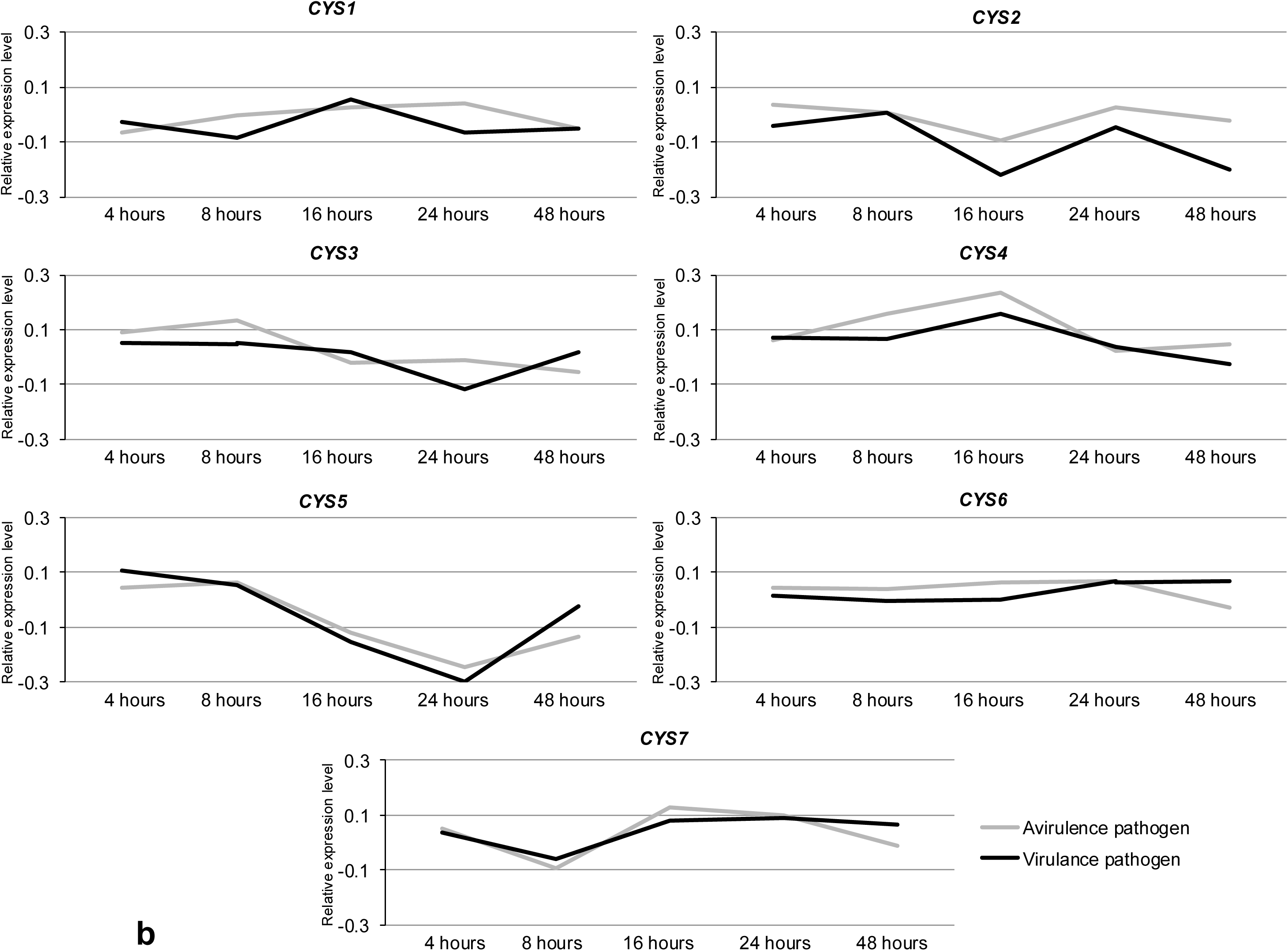
Expression profiles of seven cystatins and the five most abundant PLCPs of Arabidopsis infected with *Pseudomonas syringae* ES4326. Gene expression patterns were generated from the AtGenExpress microarray data of *Arabidopsis thaliana* (Col-0) plants challenged with *Pseudomonas syringae* strain ES4326, with or without the avirulence effector AvrRpt2 (TAIR Accession ID: 1008031517). The relative expression value was the difference between the expression value of bacteria infected treatment and mock treatment. The gray line represented the condition of bacterial infection of *Pst* ES4326 with AvrRpt2, and the black line represented the condition of bacterial infection of *Pst* ES4326. **a.** The line charts reflected the relative expression values of five PLCP genes at different time points post-bacterial infection. **b.** The line charts reflected the relative expression values of seven cystatin genes at different time points post-bacterial infection.

Although the expression levels of the five selected PLCP genes did not change significantly in response to either virulent or avirulent pathogens, RD21A was slightly downregulated and CTB3 was slightly upregulated in response to both virulent and avirulent pathogens (Figure 1a). The relative stable expression of the five most abundant PLCP genes suggested their activities might be regulated at the translational level. In contrast to the stable expression of five PLCP genes, the expression levels of seven Arabidopsis cystatin genes have relatively more significant fluctuations (Figure 1b). CYS1, CYS3, CYS6, and CYS7 did not change significantly in response to either virulent or avirulent pathogens. Therefore, these four cystatin genes may not closely relate to plant immunity to *P. syringae*. The expression of CYS2 is suppressed at 16 h and 48 h in response to the virulent *P. syringae* ES4326, while the suppression of CYS2 was attenuated in response to avirulent *P. syringae* ES4326 carrying AvrRpt2. Therefore, CYS2 and its regulated PLCPs are more likely to associate with plant basal defense. CYS4 can be specifically induced at 16 h after the inoculation of *P. syringae* ES4326 carrying AvrRpt2, suggesting that CYS4 and its related PLCPs might be associated with AvrRpt2 triggered-ETI. It is also known AvrRpt2 encodes a functional cysteine protease (Coaker, Falick et al. 2005), therefore, it will be interesting to test if CYS4 can actually inhibit the protease activity of AvrRpt2 in vivo. CYS5 was reduced two folds in response to *P. syringae* ES4326 with or without avrRpt2. Therefore, CYS5 and its related PLCPs might also participate in plant defense response.

### Co-expression Analysis of Arabidopsis PLCPs and Cystatins that May Involve in Plant Immunity

To identify if any PLCPs are co-regulated with certain cystatins in response to the infection of bacterial pathogens, we acquired the same microarray data set as described above and filtered it with 31 Arabidopsis PLCPs and seven cystatins related probes. Three PLCP genes (RDL4, RDL5 and CEP2) that have probe missing or conflicting problems (Table S1). Two housekeeping genes (UCP022280 and PTB1) were also selected in the dataset as controls. To select two housekeeping genes as controls was to avoid the accident expression of one control. The raw microarray data was processed into relative gene expression value as described in the materials and methods. The relative gene expression values of 28 PLCPs, the two control genes, and seven cystatins were used to calculate the correlation value between PLCPs and cystatins, which was presented as a heat map (Figure 2a & b). The darkness of the brick was inversely proportional to the correlation value. And the description of correlation strength was followed by the definition in the statistics textbook by Dancey et al (2004) (Dancey and Reidy 2007). Briefly, the correlation value less than 0.4 was considered as weak correlation and the correlation value greater than 0.6 was considered as strong (or significant) correlation. And the correlation value between 0.4 and 0.6 was considered as moderate correlation. At the selected significance level, none of the cystatins have significant correlation with the two housekeeping genes at the same time.

**Figure 2.**
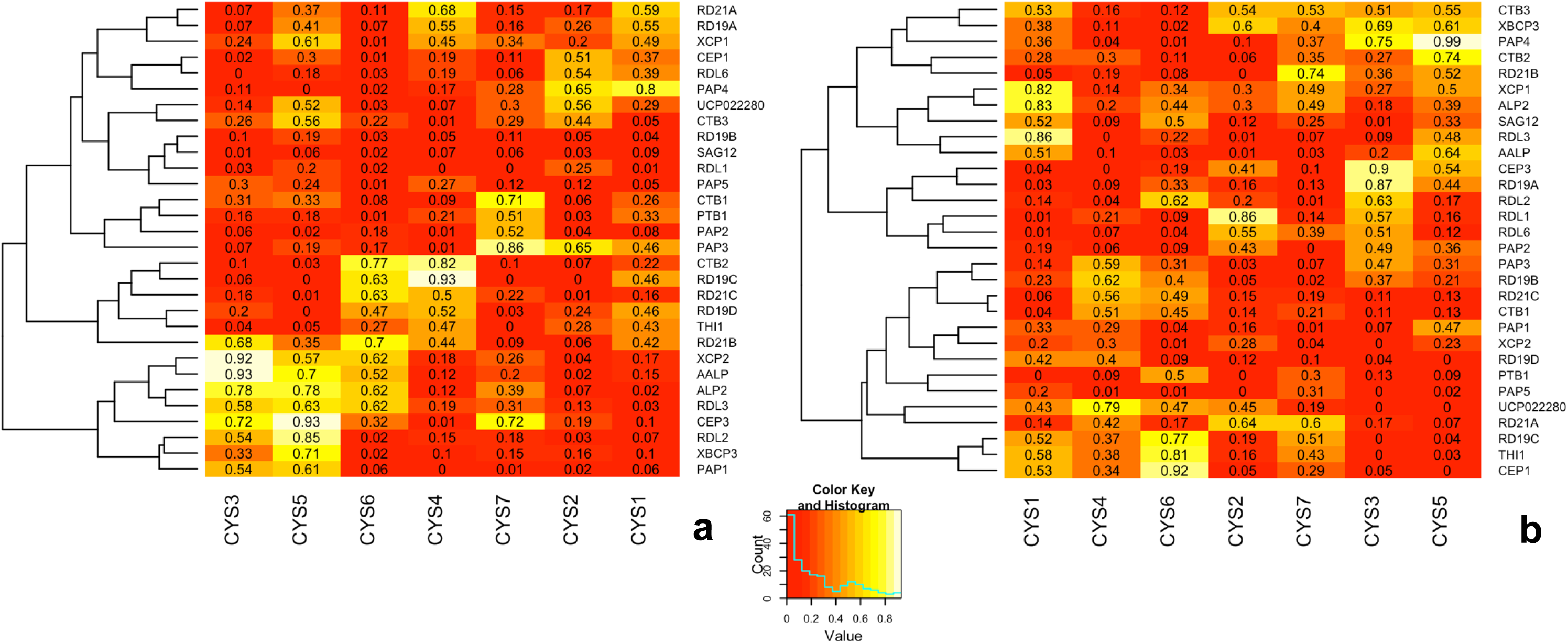
Co-expression pattern of Arabidopsis PLCPs and cystatins and clustering of PLCPs in biotic stress condition. Gene co-expression patterns were generated from the AtGenExpress microarray data of *Arabidopsis thaliana* (Col-0) plants challenged with *Pseudomonas syringae* strain ES4326, with or without the avirulence effector AvrRpt2 (TAIR Accession ID: 1008031517). The relative expression value was the difference of the expression value between bacteria infected treatment and mock treatment. The correlation value of PLCP genes (or housekeeping genes: *PTB1* and *UCP022280*) and cystatin genes were calculated based on the relative expression values of every gene at different time point post the bacterial infection. The correlation values were presented as heatmaps of co-expression pattern. The clustering of PLCP genes were based on the correlation values. **a.** The co-expression pattern of 28 PLCP genes (and two housekeeping genes) and seven cystatin genes in the condition of *Pseudomonas syringae* strain ES4326 infection were presented as the heatmap. **b.** The co-expression pattern of 28 PLCP genes (and two housekeeping genes) and seven cystatin genes in the condition of *Pseudomonas syringae* strain ES4326 with the avirulence effector AvrRpt2 infection were presented as the heatmap.

When inoculated with virulent *P. syringae* ES4326, Arabidopsis will mount a quick but weak basal defense response that will be suppressed by the virulence pathogen (Jones and Dangl 2006). Therefore, the PLCPs that have positively roles in immunity could be induced at the early stage but suppressed at the late stage of the infection with virulent bacterial pathogens. The cystatins that are co-regulated with the virulent bacterial as those PLCPs could be considered as the specific inhibitors to regulate PCLP enzyme activities at protein level. When the wild type Arabidopsis plants are inoculated with *P. syringae* ES4326 expressing AvrRpt2, the Arabidopsis R protein Rps2 recognizes AvrRpt2 and triggers ETI (Kunkel, Bent et al. 1993). The PLCPs that have positively roles in AvrRpt2-triggered ETI will be induced and their corresponding cystatins should be co-regulated to keep the balance of cysteine protease activity.

As shown in Figure 2a, when inoculated with *P. syringae* ES4326, the Arabidopsis PLCPs could be divided into five groups based on their co-expression pattern in relation to different cystatins. The top 11 PLCPs showed weak correlation with most cystatins. And four of them (RD19B, SAG12, RDL1 and PAP5) did not have significant correlation with any cystatin, that suggested they might not be related with Arabidopsis basal defense to *P. syringae*. The other 20 PLCPs showed significant correlation with at least one cystatin. And five of them (RD21B, XCP2, AALP, ALP2 and RDL3) have significant correlation value with two or three cystatins. It is possible that these five PLCPs have positive roles in Arabidopsis immunity, whose activities are suppressed by cystatins under disease susceptible conditions. Under the condition of AvrRpt2 triggered ETI (Figure 2b), the correlation pattern of PLCP-cystatin pairs did not show clear clustering. ETI is an amplified defense response, where the hypersensitive response (HR) associated PCD might regulate the expression of PLCPs and cystatins.

By comparing the correlation heat map in Figure 2a & b, XCP2 and PAP3 did not show significant correlation with any cystatin under the AvrRpt2 triggered ETI condition but showed significant correlation with two or three cystatins under susceptible condition triggered by the virulent bacteria. Thus, XCP2 and PAP3 could be classified as candidate genes that are specifically related with AvrRpt2 triggered ETI, whose activity was tightly controlled by cystatins in plant basal defense, but not under the ETI condition. The expression of CTB2, RD19C, and THI1 are strongly correlated with the expression CYS4 when Arabidopsis plants were inoculated with virulent *P. syringae* ES4326 but only has weak correlation with CYS4 under the condition of AvrRpt2 triggered ETI. Thus, we may predict CTB2, RD19C and THI1 are essential for the AvrRpt2 triggered-ETI. However, the effector AvrRpt2 itself is a cysteine protease that might affect the function of cystatins and affect the accuracy of our prediction.

### Co-expression Analysis of Arabidopsis PLCPs and Cystatins under Wounding and Drought Stress

We generated PLCP-cystatin correlation heat maps by analyzing the wounding and drought stress microarray data set (TAIR Accession ID: 1007966439 and 1007966668).

The microarray data collected from leaf tissue of Arabidopsis under the wounding stress was used to generate heat maps (Figure 3a). The PLCPs could be roughly divided into four groups based on the correlation with cystatin genes. The top 15 PLCP genes show weak or moderate correlation with all cystatins. RD21C, XCP2, RDL1 and CEP3 are strongly correlated with CYS1 or CYS3. CTB3, RDL2, CTB2 and RD19C are strongly correlated with CYS4 or CYS5. The rest PLCP genes (RD21B, THI1, PAP1, PAP5 and SAG12) are strongly correlated with CYS2 or CYS7.

**Figure 3.**
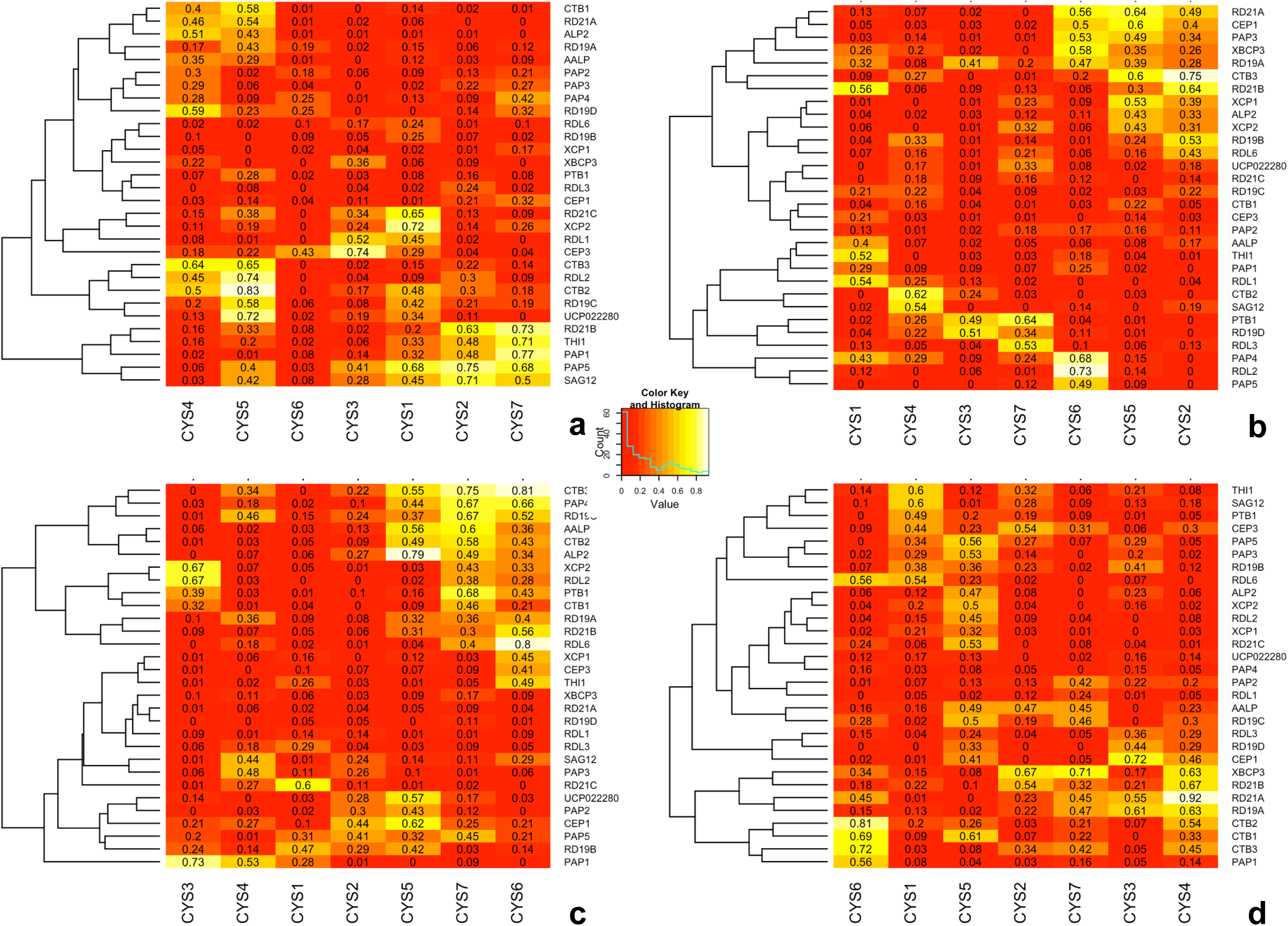
Co-expression pattern of Arabidopsis PLCPs and cystatins and clustering of PLCPs in wounding or drought stress condition. Gene co-expression patterns were generated from the AtGenExpress microarray data of *Arabidopsis thaliana* (Col-0) plants in response to wounding and drought stress (TAIR Accession ID: 1007966439 and 1007966668). The relative expression value was the difference of the expression value between bacteria infected treatment and mock treatment. The correlation value of PLCP genes (or housekeeping genes: *PTB1* and *UCP022280*) and cystatin genes were calculated based on the relative expression values of every gene at different time point post the wounding or drought stress treatment. The correlation values were presented as heatmaps of co-expression pattern. The clustering of PLCP genes were based on the correlation values. **a.** The co-expression pattern of 28 PLCP genes (and two housekeeping genes) and seven cystatin genes in Arabidopsis shoot tissue under wounding stress. **b.** The co-expression pattern of 28 PLCP genes (and two housekeeping genes) and seven cystatin genes in Arabidopsis root tissue under wounding stress. **c.** The co-expression pattern of 28 PLCP genes (and two housekeeping genes) and seven cystatin genes in Arabidopsis shoot tissue under drought stress. **d.** The co-expression pattern of 28 PLCP genes (and two housekeeping genes) and seven cystatin genes in Arabidopsis root tissue under drought stress.

The co-expression patterns of PLCP-cystatin under wounding stress (Figure 2a) is much different from the patterns under the disease condition (Figure 3a). For example, under disease conditions, the expression of XCP2, AALP, ALP2 and RDL3 are significantly correlated with at least three cystatin genes (CYS3, CYS5 and CYS6) (Figure 2a). While in response to wounding stress, these four PLCP genes are only weakly correlated with most cystatin (Figure 3a). As another example, the expression of SAG12 and PAP5 is correlated with most cystatin genes except CYS4 and CYS6 under the wounding stress, while these two PLCP genes only showed weak correlations with all seven cystatin genes in response to the inoculation of *P. syringae* ES4326.

Mechanic wounding is a mimic of damage caused by insects or animals (Jones and Dangl 2006, Habib and Fazili 2007). Under wounding stress, protease inhibitors, including cystatins, might positively regulate of plant defense by suppressing extra-plant proteases such as the digesting enzymes secreted by insects and nematodes (Habib and Fazili 2007). While PLCPs, that have roles of enhancing PCD and positively regulate defense to bacterial pathogens, could be negatively regulating plant defense to wounding or damage caused by insects (Rawlings, Tolle et al. 2004, Van der Hoorn 2008). The different co-expression patterns of PLCP-cystatins shown in Figure 2a and 3a might be consistent with this speculation.

We also analyzed the expression patterns of PLCP and cystatin genes using the microarray data collected from root tissue of Arabidopsis under the wounding stress. As shown in Figure 3b, the PLCPs could be roughly divided into three groups based on their correlation values. The top seven PLCPs showed strong correlation with CYS2, CYS5 or CYS6. The last ten PLCPs showed strong correlation with CYS1, CYS3, CYS4, CYS6 or CYS7. And the rest of PLCP genes only showed weak correlation with all cystatin genes. The clustering based on co-expression patterns suggests that some PLCPs and cystatins may serve as wounding response signaling regulators and involved in the system acquired wounding signaling transduction. The PLCP-cystatin co-expression patterns in leaf tissue (Figure 3a) is much different from the patterns in root tissue (Figure 3b). For example, six PLCP genes in Arabidopsis shoot tissue (RDL6, RD19B, XCP1, XBCP3, RDL3 and CEP1) are weakly correlated with all cystatin genes, while five different PLCPs (RD21C, RD19C, CTB1, CEP3 and PAP2) in root tissue show weak correlation with all cystatins. Therefore, Arabidopsis might have two sets of PLCPs response to wounding stress in leaf and root tissue, respectively.

Many Arabidopsis PLCPs were reported to be associated with drought tolerance, and nearly half of them were named as Responsive to Desiccation (RD) or RD-liked genes (Yamaguchi-Shinozaki, Koizumi et al. 1992). In this study, we also analyzed the co-expression patterns of PLCP-cystatin of Arabidopsis under drought conditions (Figure 3c & d). We firstly analyzed the co-expression patterns of PLCPs and cystatins in Arabidopsis shoot tissue under drought stress. As shown in Figure 3c, the PLCPs could be roughly divided into two groups based on their correlation values. The top twelve PLCP genes were significantly correlated with CYS5, CYS6 or CYS7. And the rest of PLCP genes only showed weak correlation with all cystatins. As shown in Figure 2a, when Arabidopsis plants infected with virulent bacteria, there are nine PLCP genes (RD19A, XCP1, CPE1, RDL6, CTB3, RD19B, SAG12, RDL1 and PAP5) showed weak correlation with all cystatins. And six of them (XCP1, CPE1, RD19B, SAG12, RDL1 and PAP5) show a similar co-expression pattern under the drought stress condition, where they weakly correlated with all cystatin genes. Therefore, Arabidopsis leaves response to drought stress and bacterial infection by using a similar set of PLCP and cystatin genes.

Five cystatin genes (CYS3, CYS4, CYS5, CYS6, and CYS7) showed strong correlation with most PLCP genes under the virulent bacterial infection condition, while only CYS5, CYS6 and CYS7 showed strong correlation with most PLCP genes under drought condition. Therefore, the three cystatins (CYS5, CYS6 and CYS7) might involve in regulating response to both drought stress and bacterial infection. Since both drought stress and bacterial pathogen can regulate stomatal closure, we speculate if these drought/disease shared PLCP-cystatins may have roles in regulating stomatal closure, which is worthy for further investigation in the future.

The expression of CYS2 is weakly correlated with most PLCP genes (except CEP1 and PAP5) (Figure 3c). Our previous report suggests the overexpressing of CYS2 can increase stomatal opening (Wu, 2015). Interestingly, the expression of CYS2 was suppressed under drought condition, which is usually triggering stomatal closing. Therefore, the CYS2 expression pattern is consistent with its role in regulating stomatal closing. It will be interesting to test if the CYS2 overexpression plants are also more sensitive to drought stress.

We also analyzed the co-expression patterns of PLCP and cystatin genes in Arabidopsis root tissue under drought stress, and the result was presented in Figure 3d. Comparing to the co-expression patterns of PLCP and cystatin genes in shoot tissue under drought stress (Figure 3c), the overall co-expression values of all the PLCP-cystatin were reduced. In root tissue, the expression patterns of most PLCPs, are weakly or moderately correlated with all cystatins (Figure 3d), where most PLCPs are only correlated with a few cystatins in leaf tissue (Figure 3c). Therefore, Arabidopsis might have different regulating strategies in root and shoot tissue under drought stress. Nevertheless, in root tissue, CYS5, CYS6 and CYS7 still showed strong or moderate correlation with most PLCPs (Figure 3d), which is consistent with the result in shoot tissue (Figure 3c). Therefore, these three cystatin genes (CYS5, CYS6 and CYS7) might be involved a common signaling pathway in response to drought stress. While the expression of CYS2 was slightly upregulated in root tissue, and it is significantly correlated with XBCP3, and moderately correlated with CEP, AALP, and RD21B. It is possible that CYS2 is specifically involved in regulating stomatal aperture, which is more sensitive in shoot than root tissue under the drought stress condition.

### Evaluate the Enzyme Activities of Five Arabidopsis PLCPs and the Enzyme Inhibition Ability of Seven Cystatins

In addition to the expression analysis of PLCP and cystatin genes at transcription level, we also performed protease enzyme activity and cystatin-inhibition assays to identify the interaction relationship of Arabidopsis PLCPs and cystatins at protein level.

Five abundantly expressed PLCPs (AALP, RD19A, RD19C, RD21A, and CTB3) (Figure 1b & S1) were transiently expressed in *Nicotiana benthamiana*, and crude protein extracts were used for in vitro protease activity assays following a previously described protocol (Mueller, Ziemann et al. 2013). As summarized in Figure 4, cystatins displayed distinct inhibition profiles against individual PLCPs, indicating functional diversification despite shared conserved inhibitory motifs. Specifically, CYS1 significantly inhibited CTB3, RD19A, and RD21A; CYS2 significantly inhibited AALP; CYS3 significantly inhibited RD19A and RD21A; CYS4 significantly inhibited CTB3, RD19A, and RD21A; CYS5 significantly inhibited RD19A; CYS6 significantly inhibited RD19C, CTB3, and RD19A; and CYS7 significantly inhibited RD19C and RD19A. Notably, AALP, the most abundant PLCP in Arabidopsis (Figure S1), showed limited sensitivity to cystatin inhibition, except for weak inhibition by CYS2. In contrast, RD19A was inhibited by most cystatins, consistent with the possibility that RD19A participates in more specialized stress-associated pathways that require tight regulation.

**Figure 4.**
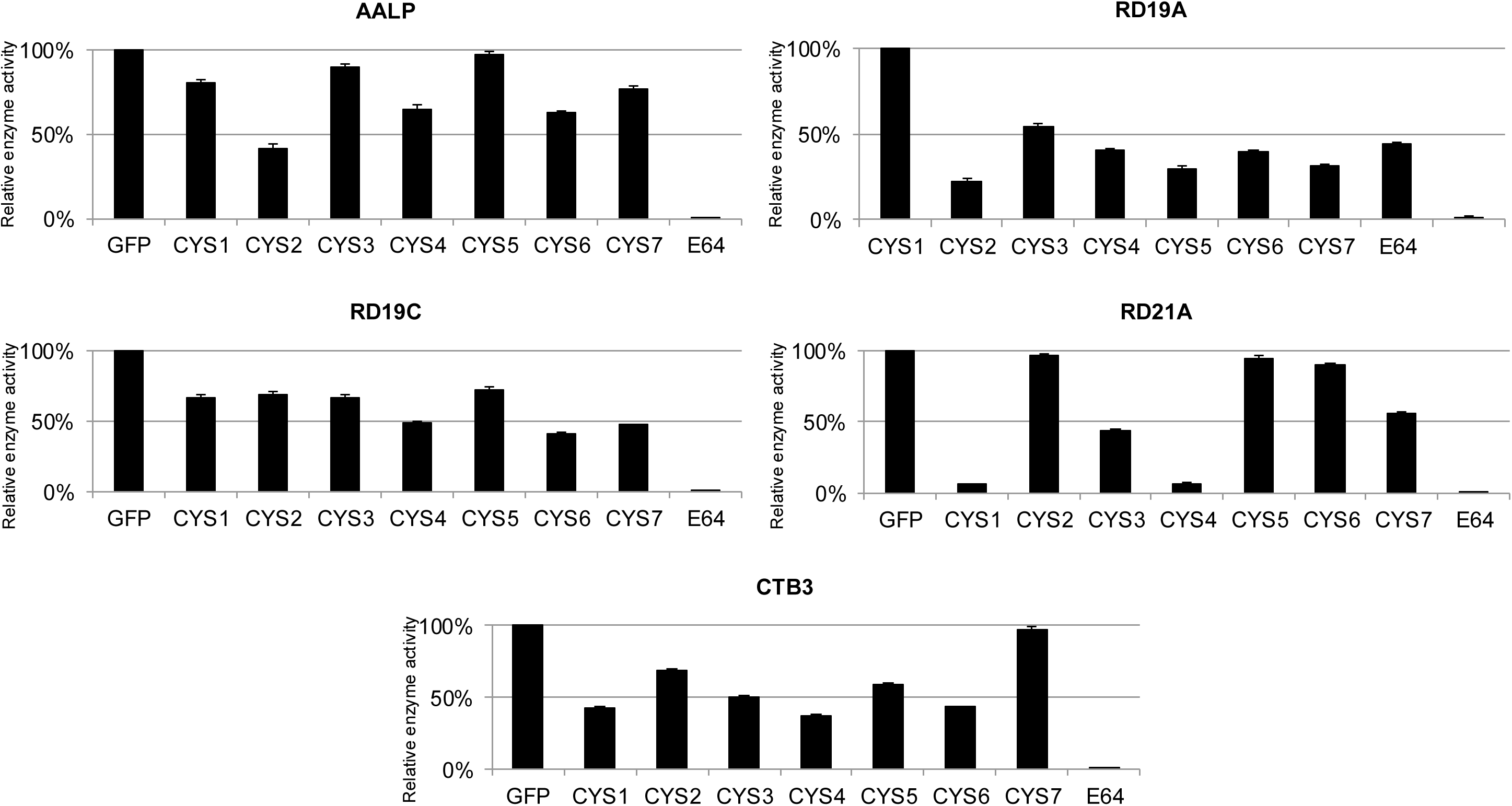
Enzyme activities of the five most abundant Arabidopsis PLCPs and the enzyme inhibition ability of seven cystatins. Five most abundant Arabidopsis PLCPs (Figure S1) were transiently expressed in *Nicotiana benthamiana* and the crude protein extract were used for enzyme activity test. The seven Arabidopsis cystatins (and the GFP control) were expressed and purified from *E. coli*-based protein expression system. The artificial cysteine protease inhibitor E64 (10µM) inhibited all five PLCPs. The means ± s.e. of enzyme activities of different PLCP and cystatin combinations were presented as the histograms. The experiment was carried out in three independent replicates.

### Prediction of Arabidopsis PLCP-Cystatin Interaction Network

We integrated PLCP-cystatin co-expression features and in vitro inhibition data to generate a hypothesis-generating PLCP-cystatin association network using the SVM implementation in the R package e1071 (Meyer and Wien 2014). For each PLCP-cystatin pair, co-expression correlations from six conditions (virulent infection, avirulent infection, wound_shoot, wound_root, drought_shoot, and drought_root) were used as features. Inhibition outcomes from the in vitro assays were used to label the subset of tested PLCP-cystatin pairs as ‘associated’ or ‘not associated’ for model training. Model parameters (cost and gamma) were selected using standard k-fold cross-validation (Hsu, Chang et al. 2003). Given the limited size and scope of the training set, the SVM model was used as an integrative scoring framework rather than as a definitive predictor of physical interaction. The full set of PLCP-cystatin scores was then visualized as a network in Cytoscape (Figure 5a).

**Figure 5.**
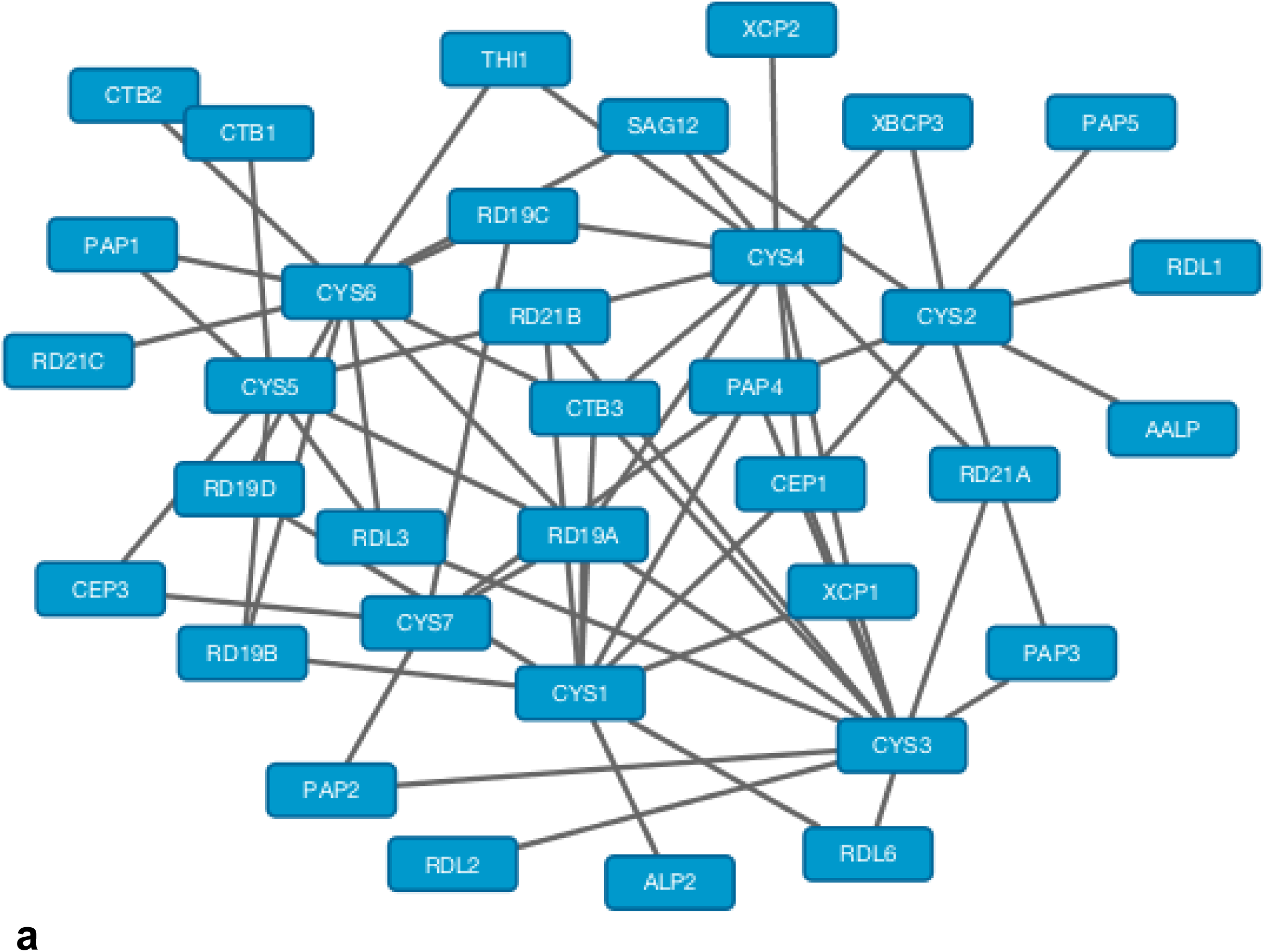
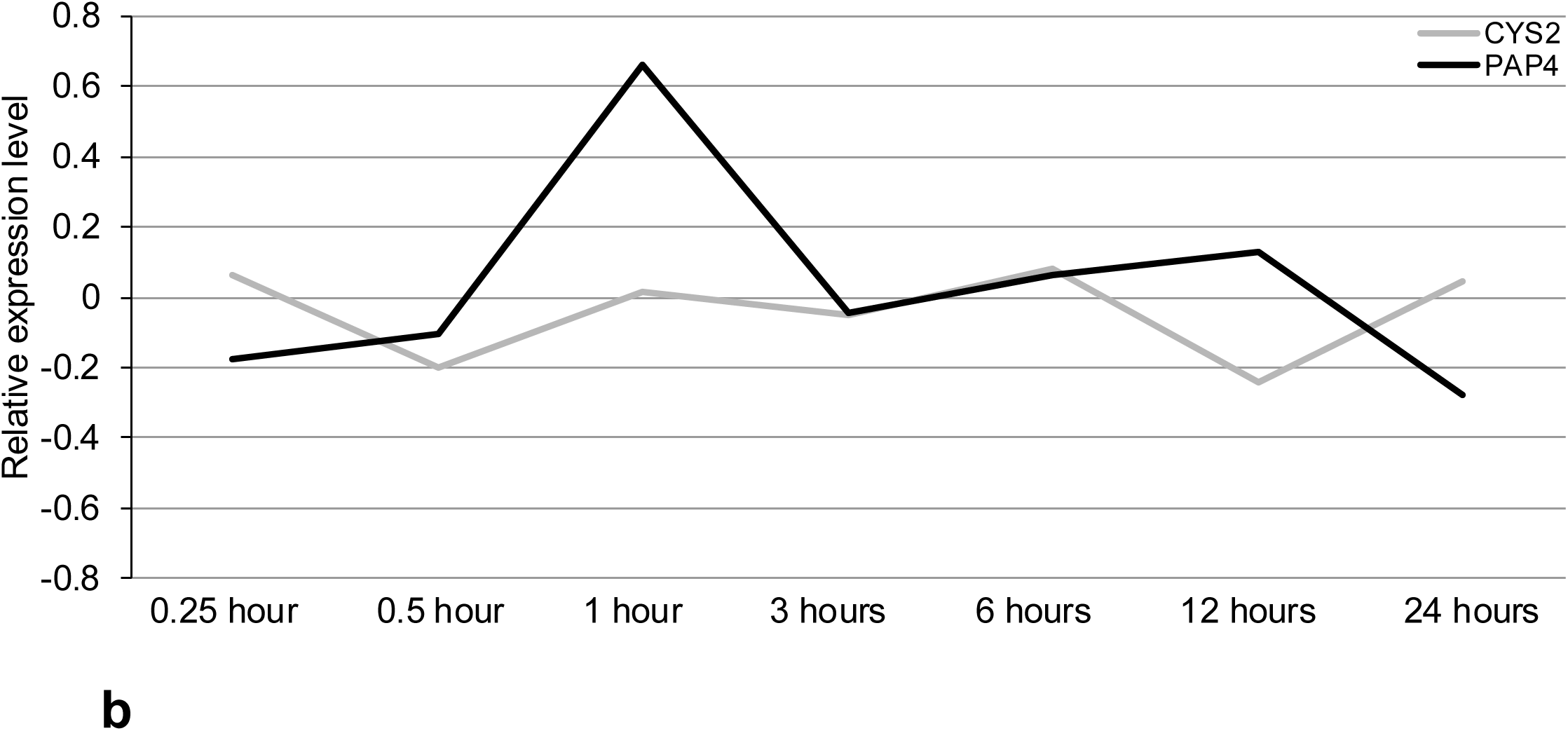
PLCP-cystatin interaction network and the expression profiles of *CYS2* and *PAP4* under drought stress. **a.** The interaction network of 28 PLCP genes and seven cystatin genes were generated based on the prediction result in Table S3, using Cytospace program. **b.** Gene expression patterns of *CYS2* and *PAP4* were generated from the AtGenExpress microarray data of *Arabidopsis thaliana* (Col-0) plants in response to drought stress (TAIR Accession ID: 1007966668). The line chart represented the relative expression value of *CYS2* and *PAP4* at different time points post the drought stress treatment.

As shown in Figure 5a, CYS3 interacts with at least 12 most PLCPs (RD21A, RD21B, RDL2, RDL3, RDL6, CEP1, XCP1, PAP2, PAP3, PAP4, RD19A and CTB3), while CYS7 interacts with only five PLCPs (CEP3, PAP2, PAP4, RD19A and RD19C). Ten PLCPs interact with at least three cystatins (RD21A, RD21B, RDL3, CEP1, XCP1, PAP4, RD19A, RD19B, RD19C, and CTB3). Therefore, these ten PLCPs may have important functions in Arabidopsis, where their activities are tightly regulated by several different cystatins. Interestingly, in the RD19 family (Table S1), all three family members belonged to this group, which suggested the RD19 family was conserved and involved in regulating some essential biological activities in Arabidopsis. In the 28 PLCPs, only 8 of them (RD21C, RDL1, RDL2, PAP2, PAP5, AALP, ALP2, CTB1 and CTB2) are predicted to be related with only one cystatin. All the others are predicted to be related with two or more cystatins. That indicated most PLCPs and cystatins may have functional redundancy and synergy.

The PLCP-cystatin interaction network provides a general view of the PLCP-cystatin interplay. When characterize biological function of a particular PLCP or cystatin gene, we may consider analyzing the co-related genes as a group in considering of gene redundancy problem. For example, in our previous study, the effector AvrRxo1 from *Xanthomonas oryzae* pv. *oryzicola* (*Xoc*) was identified to be able specifically induce the expression of Arabidopsis CYS2 (Wu, 2015). Overexpression of CYS2 increases stomata aperture size and suppresses plant immunity. As shown in Figure 2a and 3c, the expression of CYS2 had strong correlation with PAP4 when plants were infected with bacterial pathogen. However, the correlation is not significant when plants were treated with drought stress. As shown in Figure 5b, expression of CYS2 was suppressed at the early stage of drought treatment, while the expression PAP4 was significantly induced. Considering the overexpression of CYS2 can increase stomata aperture size, we predict the overexpression or induced expression of PAP4 may reduce the stomatal aperture size, which can prevent water loss as early response to drought stress.

### Conclusions and Perspectives

In this study, we analyzed the microarray data of Arabidopsis plant in response to bacterial pathogen *P. syringae* ES4326, wounding, and drought stress, and predicted the relationship between 28 PLCP and seven cystatin genes in Arabidopsis. The heat map of gene co-expression summarizes the correlation between different PLCP and cystatin genes, which provides an overview for the interplay of PLCP-cystatin. In general, the expression of some cystatin genes is regulated by *P. syringae* ES4326, which have roles to regulate PLCP enzyme activity. Both PLCPs and cystatins have variable expression in different tissues (root vs shoot). By comparing the correlation patterns of PLCPs and cystatins under different conditions, we predicted several candidate PLCPs and cystatins, which may specifically involve in a particular signal transduction pathway in response to abiotic or biotic stress.

In the future, to improve the interaction network, we need analyze more microarray (or RNA-seq) data, including other effector-triggered immunity or other types of pathogen infection. It is known, many PLCPs are only expressed in seed or reproduction organs. Therefore, expression data from more tissue types are needed to be analyzed in the future. In addition to analysis of Arabidopsis PLCP and cystatin genes at the transcriptional level, it will be valuable to perform more comprehensive enzyme activity and inhibition assays between seven Arabidopsis cystatins and all thirty-one PLCPs at the protein levels. A comprehensive yeast two-hybrid assay or co-immunoprecipitation to validate the physical interactions between the PLCPs and cystatins can also provide insights about the interplay of PLCP-Cystatin. All this information will help us build more comprehensive interaction network of the PLCP-cystatin that will be useful for studying plant immunity and resistance to abiotic stress.

## Material and Methods

### Acquiring the raw microarray data

The microarray data was downloaded from Atgenexpress database on TAIR website (https://www.arabidopsis.org). Three data sets used in this study were: (1) “Pseudomonas half leaf injection” (TAIR Accession ID: 1008031517) (Wang, Weaver et al. 2005); (2) “Wounding stress time course” (TAIR Accession ID: 1007966439) (Kilian, Whitehead et al. 2007); (3) “Drought stress time course” (TAIR Accession ID: 1007966668) (Kilian, Whitehead et al. 2007).

### Microarray data processing

The raw microarray data was separated by treatments and time points, and organized into six data sets: (1) virulent bacteria challenge, (2) avirulent bacteria challenge, (3) wounding stress in shoot tissue, (4) wounding stress in root tissue, (5) drought stress in shoot tissue, and (6) drought stress in root tissue. We then acquired the information for 28 PLCP genes, seven cystatin genes and two housekeeping genes (Table S1). Although 31 putative PLCP genes were predicted in Arabidopsis genome, three of them (RDL4, RDL5, and CEP2) have problems of missing or conflicting probes in the microarray data sets, therefore, they were excluded in this study. The averaged microarray signal values of three replicates were transformed into relative expression values by using the formula: Relative expression value = log10 (averaged microarray signal value of the experimental treatment) -log10 (averaged microarray signal value of the mock treatment). The relative expression values were used to calculate the correlation value between each pair of PLCP and cystatin, or the two housekeeping genes. The R software (Version 3.2) with the package of “heatmap.plus” was used for producing the heat maps (Day 2012).

### Cloning of the PLCPs and cystatin genes

Five Arabidopsis PLCP genes (AALP, RD19A, RD19C, RD21A and CTB3) and seven Arabidopsis cystatin genes (CYS1, CYS2, CYS4, CYS5, CYS6, and CYS7) were PCR amplified from the cDNAs of Arabidopsis Col-0 using primers list in Table S2. The Cys3 was failed to be amplified from cDNAs, and it was synthesized by GenScript (Piscataway, NJ). All full length PLCP and CYS genes were cloned into pDonr207 vector (Invitrogen, Carlsbad, CA) and the correct sequence was confirmed by sequencing as the core facility of Virginia Bioinformatics Institute (Blacksburg, VA). The GFP gene was amplified from p519gfp (Matthysse, Stretton et al. 1996). All cloned genes were subcloned into the Gateway compatible binary vector pEarleygate 102 (Earley, Haag et al. 2006) through LR® cloning following the manufacturer’s instruction (Invitrogen). Seven cystatin and the GFP genes were LR® cloned into a Gateway compatible expression vector pGEX4T-Des vector (Han et al, 2015).

### Protein expression and purification of cystatins

The expression plasmids were transformed into *E. coli* C41 cells (Lucigen, Middleton, WI) and grown overnight at 37 °C in 50 ml of LB medium supplemented with 100 mg/L carbenicillin. The bacteria culture was sub-cultured into 1 liter of LB medium containing 100 mg/L carbenicillin till to the OD600 is about 0.8 unit. Isopropyl-β-D-thiogalactoside (IPTG) was added to a final concentration 0.5 mM with subsequent incubation at 28°C/220 rpm for 8 hours. The bacterial cells were harvested by centrifugation at 5,000 rpm for 10 min at 4°C and the bacteria pellet was resuspended in 50 ml of binding buffer (50 mM Tris buffer, pH 7.5, 500 mM NaCl, 2 mM 2-mercaptoethanol, 1mM PMSF). The bacterial cells were then broken by incubating them with 1 µg/ml lysozyme on ice for one hour. Afterward, they were subjected to three rounds of sonication on ice at 50% duty for 8 seconds each time. The resulting lysate was centrifuged at 12,000 g for 20 minutes at 4°C, and the supernatant was collected and used for further purification. Glutathione resin (GenScript) was equilibrated with binding buffer and added to the cleared supernatants. The mixture was then incubated at 4°C for one hour, subsequently loaded onto a 5 ml column (1 cm x 20 cm) (Bio-Rad, Hercules, CA). The column was washed three times with 50 ml of washing buffer (50 mM Tris-HCl buffer, pH 8.0, 300 mM NaCl, 1 mM DTT) to remove any nonspecifically bound proteins. The bound proteins were then eluted with 10 x 1 ml of washing buffer containing 10 mM glutathione. The protein concentration was measured using NanoDrop® (Thermo Scientific, Hudson, NH), and the protein purity was determined by running an SDS-PAGE gel and visualizing it with Coomassie blue staining.

### Agrobacterium-mediated transient assay and in vitro protease activity assay

To achieve in planta post-translational activation of the PLCPs, the five PLCP genes were transiently expressed in *N. benthamiana* following the protocol described previously (Zhao, Dahlbeck et al. 2011). Briefly, the binary constructs were transformed into Agrobacterium strain GV2260 by electroporation (Traore and Zhao 2011). The Agrobacterium cells were resuspended in 10 mM MgCl2 and adjusted to OD600 = 0.4 units and infiltrated into the leaf of *N. benthamiana* using a blunt end syringe without needle. The infiltrated plants were incubated at room temperature under continuous light. Leaf disks were collected at four days post infiltration using a cork borer (size 3) (Starkey and Rahme 2009, Mueller, Ziemann et al. 2013). Three leaf disks of each sample were pooled for protein extraction. The crude protein extract was used for in vitro protease activity assay as described by Mueller et al (Mueller, Ziemann et al. 2013). Briefly, the leaf discs were grounded in 100 µL reaction buffer (50 mM sodium phosphate, pH 6.0, 600 mM NaCl, 4 mM EDTA, 2 mM DTT) on ice and centrifuged at 12,000 rpm for 5 minutes at 4°C. Forty µL of supernatant as the PLCP extracts was aliquot into two micro-tubes as the two replicates. The PLCP extracts were either added with 10 µM GFP or cystatin proteins. As control, 5 mM of E64 (the cysteine protease inhibitor) (Sigma, St Louis, MI) was added to one set of PLCP samples. The reaction mixture was incubated at room temperature for 10 minutes, followed by adding 10 µL of 10 mM of Z-Phe-Arg-AMC (Sigma, St Louis, MI) as the protease substrate. The mixture was further incubated at room temperature for another 10 minutes. The reduction of fluorescence signal as the reflection of PLCP enzyme activity was measured using a Microplate Reader (BioTeck®, Synergy HT Multi-Mode). The values of fluorescence signals were used to compare the difference between PLCP samples incubated with either GFP or Cystatins, or E64.

### Prediction of the PLCP-cystatin interaction network

The SVM (support vector machines) package (’e1071’) in R software (Version 3.2) were used for modelling the PLCP-cystatin interaction network (Meyer and Wien 2014). Firstly, we separated the PLCP-cystatin co-expression correlation patterns into either biotic stress or abiotic stress conditions and grouped them into a six-dimension data set. The dimensions were (1) “virulence, (2) avirulence”, (3) “wound_shoot”, (4) “wound_root”, (5) “drought_shoot”, and (6) “drought_root”. Each pair of PLCP-cystatin could be considered as one data point in the six-dimension data matrix, and the SVMs can provide a proper binary classification model to divide all the points into two groups (either “related” or “nonrelated”). Secondly, we predicted the PLCP-cystatin interaction pairs based on in vitro enzyme activity assay as described previously. The cystatins that suppressed more than 50% reduction of the fluorescence signals were considered as the significant inhibitors to the corresponding PLCPs, and these pairs of cystatins and PLCPs were considered to be co-related to each other. We simplified the training input by considering the cystatin that reduced more than 50% PLCP enzyme activity as the inhibitor of this PLCP and marked this pair of PLCP-cystatin interaction as “1”. And the unqualified pairs were marked as “0”, to represent no co-relation. The 35 pairs (five PLCPs and seven cystatins) of PLCP-cystatin interaction values (“1” or “0”) and their co-expression correlation values were used as the training data to select parameters for the SVMs models (Table S3). The parameter was selected following the standard protocol of k-fold cross-validation (Hsu, Chang et al. 2003). We chose the parameter setting as: cost = 5, gamma = 1, which had the highest score ranking in the cross-validation, and it provided 100% accuracy on classifying the training data set. Finally, we processed the binary classification on the total data set of PLCP-cystatin co-expression correlations, which integrated all the six sets of PLCP-cystatin co-expression data we used in the former analysis. And the raw output of the SVMs modeling was used to generate the interaction network using the Cytoscape (http://www.cytoscape.org/index.html).

## Acknowledgement

Financial support was provided by grants from US National Science Foundation (NSF) (IOS-0845283 to BZ), and the Virginia Agricultural Experiment Station (VA-160144). This project was also funded, in part, with an integrated, internal competitive grant from VAES, VCE, and the College of Agriculture and Life Sciences at Virginia Tech.

---

**Figure S1.**
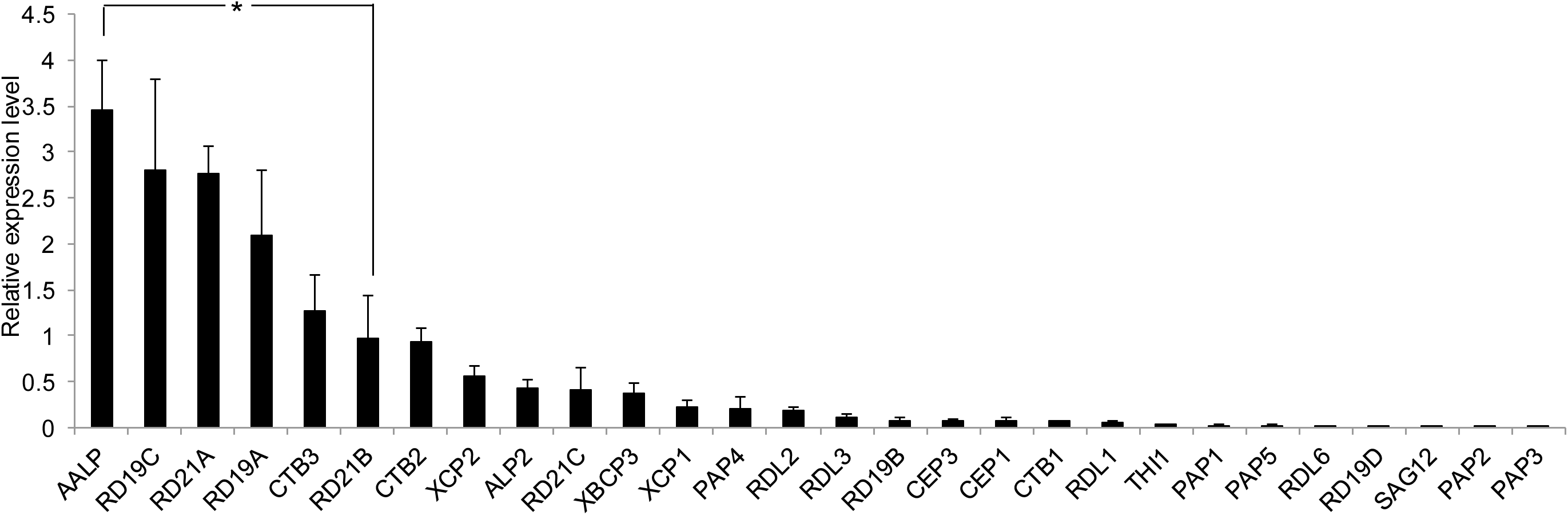
Expression levels of 28 PLCP genes in Arabidopsis shoot and root tissues. The expression values of 28 PLCP genes (Table S1) in Arabidopsis were generated from the AtGenExpress microarray data of *Arabidopsis thaliana* (Col-0) plants challenged with *Pseudomonas syringae* strain ES4326 (TAIR Accession ID: 1008031517), and the wounding and drought stress microarray data set (TAIR Accession ID: 1007966439 and 1007966668). The means ± s.e. of expression values of 28 PLCP genes in “non-treatment” conditions of all micro array data sets were presented. The asterisk represented significant difference (p-value< 0.05, Student’s t-test).

**Table S1.**
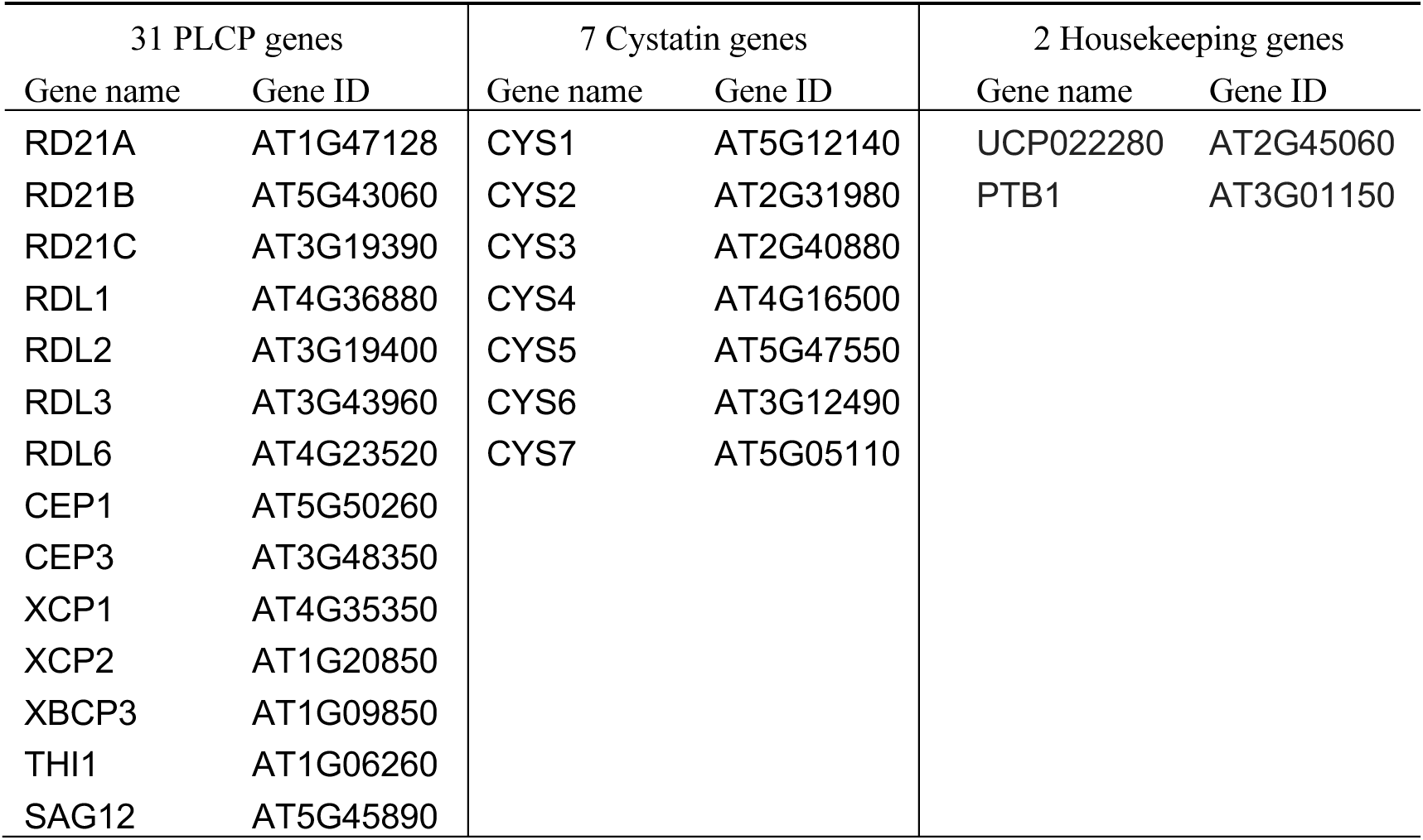
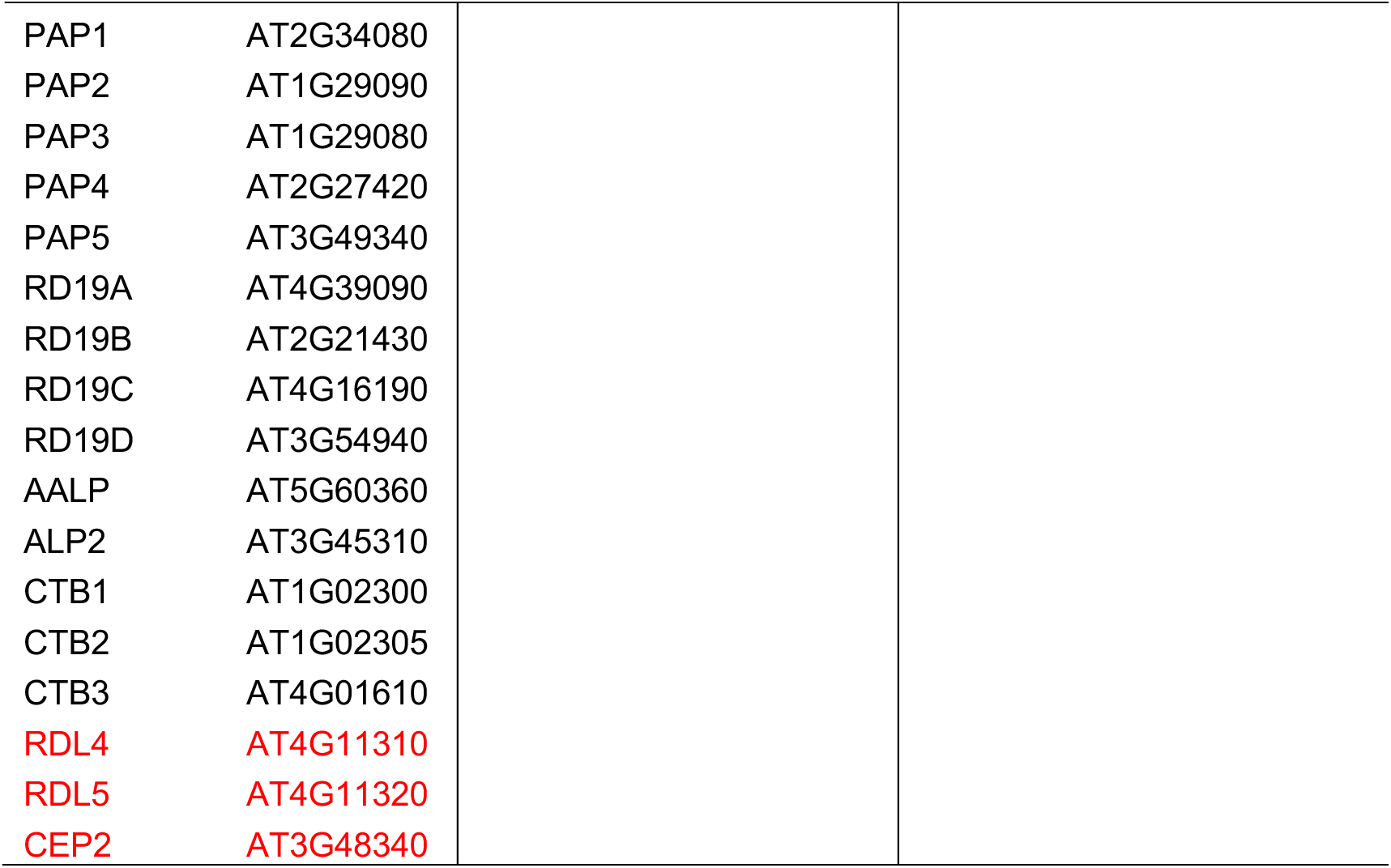
Gene names and TAIR IDs for PLCP, cystatin and housekeeping genes used in this study. The three PLCP genes have probe missing or conflicting problem are labeled in red color.

| 31 PLCP genes |  | 7 Cystatin genes |  | 2 Housekeeping genes |  |
| --- | --- | --- | --- | --- | --- |
| Gene name | Gene ID | Gene name | Gene ID | Gene name | Gene ID |
| RD21A | AT1G47128 | CYS1 | AT5G12140 | UCP022280 | AT2G45060 |
| RD21B | AT5G43060 | CYS2 | AT2G31980 | PTB1 | AT3G01150 |
| RD21C | AT3G19390 | CYS3 | AT2G40880 |  |  |
| RDL1 | AT4G36880 | CYS4 | AT4G16500 |  |  |
| RDL2 | AT3G19400 | CYS5 | AT5G47550 |  |  |
| RDL3 | AT3G43960 | CYS6 | AT3G12490 |  |  |
| RDL6 | AT4G23520 | CYS7 | AT5G05110 |  |  |
| CEP1 | AT5G50260 |  |  |  |  |
| CEP3 | AT3G48350 |  |  |  |  |
| XCP1 | AT4G35350 |  |  |  |  |
| XCP2 | AT1G20850 |  |  |  |  |
| XBCP3 | AT1G09850 |  |  |  |  |
| THI1 | AT1G06260 |  |  |  |  |
| SAG12 | AT5G45890 |  |  |  |  |

|  |  |
| --- | --- |
| PAP1 | AT2G34080 |
| PAP2 | AT1G29090 |
| PAP3 | AT1G29080 |
| PAP4 | AT2G27420 |
| PAP5 | AT3G49340 |
| RD19A | AT4G39090 |
| RD19B | AT2G21430 |
| RD19C | AT4G16190 |
| RD19D | AT3G54940 |
| AALP | AT5G60360 |
| ALP2 | AT3G45310 |
| CTB1 | AT1G02300 |
| CTB2 | AT1G02305 |
| CTB3 | AT4G01610 |
| RDL4 | AT4G11310 |
| RDL5 | AT4G11320 |
| CEP2 | AT3G48340 |

**Table S2.**
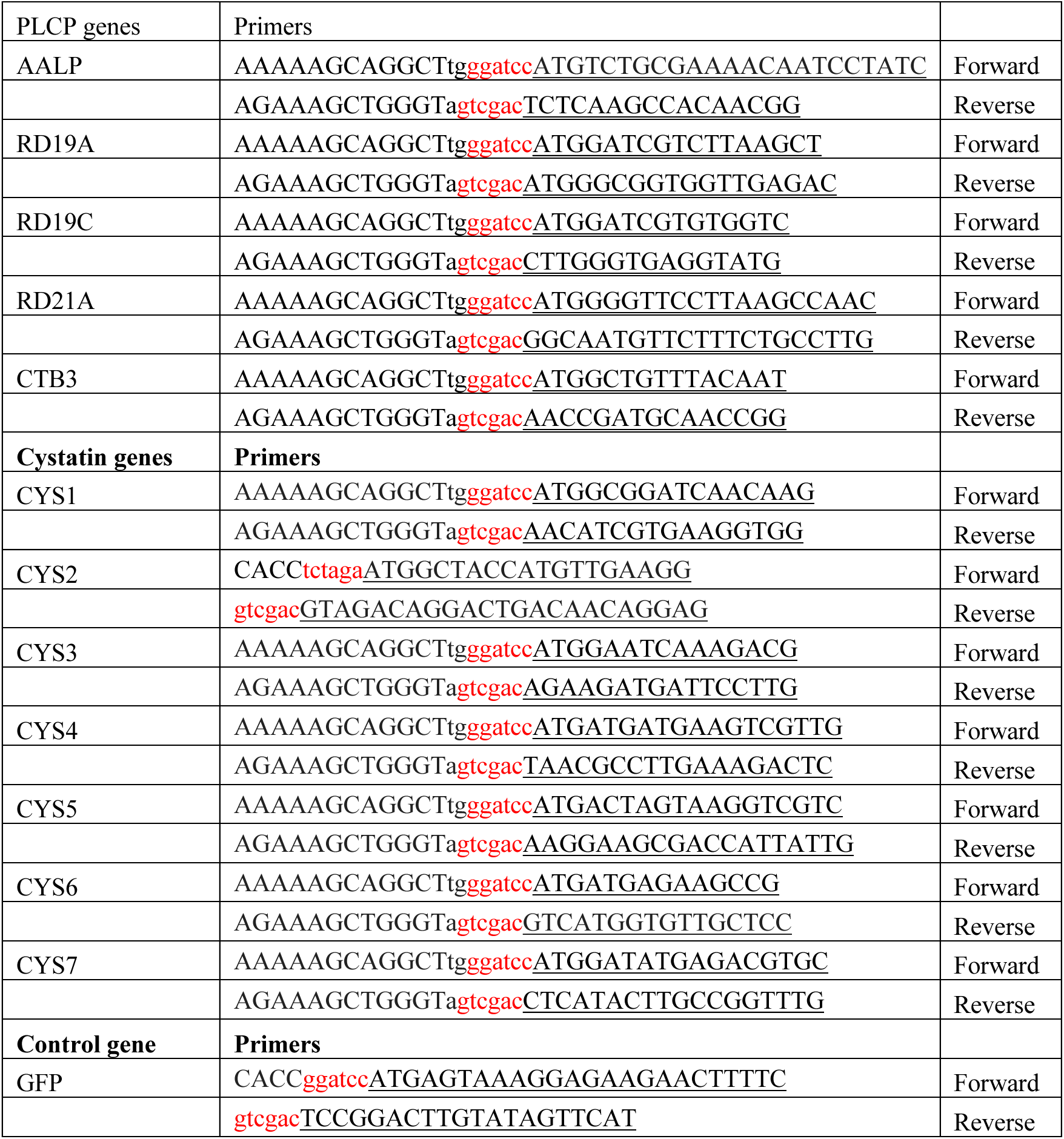
Sequences of primers used in this study. Restriction sites are labeled in red color, and the sequences from cDNA are underlined.

**Table S3.** Predicted relationship between seven cystatins and 28 PLCPs. The predicted relationship between seven cystatin genes and 28 PLCP genes (and two housekeeping control genes) were generated by SVMs modeling with the R package “e1071”. As presented in the table, “0” represented “non-related” and “1” represented “related”.

| PLCP/ control<br>genes | Cystatin genes |  |  |  |  |  |  |
| --- | --- | --- | --- | --- | --- | --- | --- |
|  | CYS1 | CYS2 | CYS3 | CYS4 | CYS5 | CYS6 | CYS7 |
| PTB1 | 0 | 0 | 0 | 0 | 0 | 0 | 0 |
| UCP022280 | 0 | 0 | 0 | 0 | 0 | 0 | 0 |
| RD21A | 1 | 0 | 1 | 1 | 0 | 0 | 0 |
| RD21B | 1 | 0 | 1 | 1 | 1 | 0 | 0 |
| RD21C | 0 | 0 | 0 | 0 | 0 | 1 | 0 |
| RDL1 | 0 | 1 | 0 | 0 | 0 | 0 | 0 |
| RDL2 | 0 | 0 | 1 | 0 | 0 | 0 | 0 |
| RDL3 | 0 | 0 | 1 | 0 | 1 | 1 | 0 |
| RDL6 | 1 | 0 | 1 | 0 | 0 | 0 | 0 |
| CEP1 | 1 | 1 | 1 | 1 | 0 | 0 | 0 |
| CEP3 | 0 | 0 | 0 | 0 | 1 | 0 | 1 |
| XCP1 | 1 | 0 | 1 | 1 | 0 | 0 | 0 |
| XCP2 | 0 | 0 | 0 | 1 | 0 | 0 | 0 |
| XBCP3 | 0 | 1 | 0 | 1 | 0 | 0 | 0 |
| THI1 | 0 | 0 | 0 | 1 | 0 | 1 | 0 |
| SAG12 | 0 | 1 | 0 | 1 | 0 | 1 | 0 |
| PAP1 | 0 | 0 | 0 | 0 | 1 | 1 | 0 |
| PAP2 | 0 | 0 | 1 | 0 | 0 | 0 | 1 |
| PAP3 | 0 | 1 | 1 | 0 | 0 | 0 | 0 |
| PAP4 | 1 | 1 | 1 | 0 | 0 | 0 | 1 |
| PAP5 | 0 | 1 | 0 | 0 | 0 | 0 | 0 |
| RD19A | 1 | 0 | 1 | 1 | 1 | 1 | 1 |
| RD19B | 1 | 0 | 0 | 0 | 1 | 1 | 0 |
| RD19C | 0 | 0 | 0 | 1 | 0 | 1 | 1 |
| RD19D | 1 | 0 | 0 | 0 | 0 | 1 | 0 |
| AALP | 0 | 1 | 0 | 0 | 0 | 0 | 0 |
| ALP2 | 1 | 0 | 0 | 0 | 0 | 0 | 0 |
| CTB1 | 0 | 0 | 0 | 0 | 1 | 0 | 0 |
| CTB2 | 0 | 0 | 0 | 0 | 0 | 1 | 0 |
| CTB3 | 1 | 0 | 1 | 1 | 0 | 1 | 0 |

